# A novel LC system embeds analytes in pre-formed gradients for rapid, ultra-robust proteomics

**DOI:** 10.1101/323048

**Authors:** Nicolai Bache, Philipp E. Geyer, Dorte B. Bekker-Jensen, Ole Hoerning, Lasse Falkenby, Peter V. Treit, Sophia Doll, Igor Paron, Florian Meier, Jesper V. Olsen, Ole Vorm, Matthias Mann

## Abstract

**To further integrate mass spectrometry (MS)-based proteomics into biomedical research and especially into clinical settings, high throughput and robustness are essential requirements. They are largely met in high-flow rate chromatographic systems for small molecules but these are not sufficiently sensitive for proteomics applications. Here we describe a new concept that delivers on these requirements while maintaining the sensitivity of current nano-flow LC systems. Low-pressure pumps elute the sample from a disposable trap column, simultaneously forming a chromatographic gradient that is stored in a long storage loop. An auxiliary gradient creates an offset, ensuring the re-focusing of the peptides before the separation on the analytical column by a single high-pressure pump. This simplified design enables robust operation over thousands of sample injections. Furthermore, the steps between injections are performed in parallel, reducing overhead time to a few minutes and allowing analysis of more than 200 samples per day. From fractionated HeLa cell lysates, deep proteomes covering more than 130,000 sequence unique peptides and close to 10,000 proteins were rapidly acquired. Using this data as a library, we demonstrate quantitation of 5200 proteins in only 21 min. Thus, the new system-termed Evosep One-analyzes samples in an extremely robust and high throughput manner, without sacrificing in depth proteomics coverage.**

---

Bottom-up proteomics is a highly successful and generic technology, which now allows the analysis of complex samples ranging from bacteria through cell line systems and even human tissue samples (Aebersold and Mann, 2016). State-of-the-art workflows begin with a robust sample preparation to digest proteins and harvest purified peptides (Kulak et al., 2014), which are separated by a liquid chromatography (LC) system before they are analysed by a mass spectrometer (MS). Established software solutions automatically interpret the acquired spectra, generating lists of thousands of quantified proteins (Bekker-Jensen et al., 2017; Bruderer et al., 2017; Cox and Mann, 2008; Kelstrup et al., 2018; Kulak et al., 2017; Meier et al., 2018).

The current performance level is a result of improvements not only in the mass spectrometric components both also the chromatographic part of the LC-MS workflow. In the quest for ever increasing chromatographic separation power, columns have become longer and particle sizes smaller - now reaching the sub 2 μm range. This may require pump pressures in excess of 1000 bar, presenting great engineering challenges for both the pumps and the entire LC system, often limiting robustness in routine operation. Thus, chromatography remains a weak link in MS-based proteomics workflows, leading to calls for new approaches (Riley et al., 2016). Furthermore, irreproducibility of retention times within and between laboratories severely limits strategies that rely on the transfer of accurate retention times, especially targeted proteomics (Picotti and Aebersold, 2012), data independent acquisition (Gillet et al., 2016) and ‘match between runs’ at the MS level (Cox et al., 2014; Geiger et al., 2012).

There is great interest in applying the increasing power of MS-based proteomics to diagnostic and clinical questions (Geyer et al. 2017). ‘Clinical proteomics’, however, requires far more stability and reproducibility than that available even in the most advanced MS-based proteomics laboratories. Note that irreproducibility and robustness issues are not features of LC-MS *per se*, as the measurement of small molecules is firmly established in clinical laboratories around the world, which routinely measure hundreds of samples per day. The two key differences of these LC systems to the one applied in proteomics are their much larger column diameters (20-fold) and flow rates (1000-fold), making them much easier to control and less error-prone. Increasing the flow rates to achieve greater robustness has already been advocated in the context of cancer proteomics (Liu et al., 2013). However, the signal intensity in electrospray ionization is concentration dependent and reducing sensitivity at higher flow rates, which limits these approaches to a few μl/min. Apart from high robustness, throughput is the other central requirement for MS-based proteomics, if it is to enter routine clinical use. Instead, current proteomics workflows generally employ fractionation - multiplying measurement time - or use relatively long gradient times.

In a recent large-scale plasma proteomics study measured in our laboratory, involving more than a thousand samples, 80% of the overall down time was attributable to the HPLC system rather than the MS. At the same time, column equilibration, loading and washing steps between runs limited the attractiveness of very short gradients (Geyer et al. 2016a; Geyer et al. 2016b).

A number of years ago, some of the current authors devised a very different sample loading and injection approach. Termed speLC, for solid-phase-extraction (nano) liquid chromatography, it was intended for very high sample throughput needed for clinical application (Falkenby et al., 2014). The speLC made use of the same StageTips that are commonly employed in proteomics for micro-scale purification of peptides and crude manual fractionation (Ishihama et al., 2006; Rappsilber et al., 2003; Wisniewski et al., 2013). Instead of eluting into the autosampler vial of the HPLC system, a low-pressure pump passed a 5-10 min gradient through the StageTip itself and directly towards the MS. The speLC system was capable of analysing 192 *E.coli* samples in only 30 h, as well as identifying more than 500 proteins from a HeLa cell lysate in less than 10 min (Falkenby et al., 2014). In subsequent work, speLC was combined with pre-fractionation such as 1D gel electrophoresis or strong cation exchange (SCX), capitalizing on its ability to analyse each of the fractions in 10 min or less (Binai et al., 2015). Although useful for simple protein mixtures, the low-pressure elution from StageTips and use of only very short analytical columns inherently limited chromatographic separation power of this system.

In the work reported here, we aimed to preserve the benefits of the original speLC device while also achieving the desirable features of modern HPLC instruments. We realized this goal by coupling elution through the StageTips to a novel downstream workflow. In the Evosep One design, peptides are eluted at low pressure and flow rates of tens of μl/min from a special StageTip - termed EvoTip™. Notably, the gradient along with the eluted analytes are captured in a long capillary loop. A single high-pressure pump then applies the stored gradient to an analytical nano-scale column. This results in undiminished chromatographic separation performance while eliminating the need to form a gradient at high pressure. Thus, this layout marries the convenience and robustness of large columns, high-flow systems with the sensitivity of narrow column diameters and low-flow rates of nano-LC systems. We further detail the principle of operation and development of the Evosep instrument in detail and investigate its robustness, throughput, and reproducibility in typical applications encountered in MS-based proteomics.

## EXPERIMENTAL PROCEDURES

*Description of the liquid chromatography system* - The Evosep One incorporates four low-pressure single stroke piston pumps (A, B, C and D) and one high-pressure single stroke piston pump (HP) (Fig. 1). Together they create a separate low-and high-pressure sub-system. Each pump is equipped with a pressure (PSx) and flow sensor (FSx) to monitor and precisely control the flow of the individual solvent. A custom 12-port valve (V12) diverts the flow of the low-pressure pumps either towards the solvent bottles (Sx) for refilling or towards the system for analysis. The high-pressure pump has a separate 6-port valve (V6) for refilling.

**Fig. 1:**
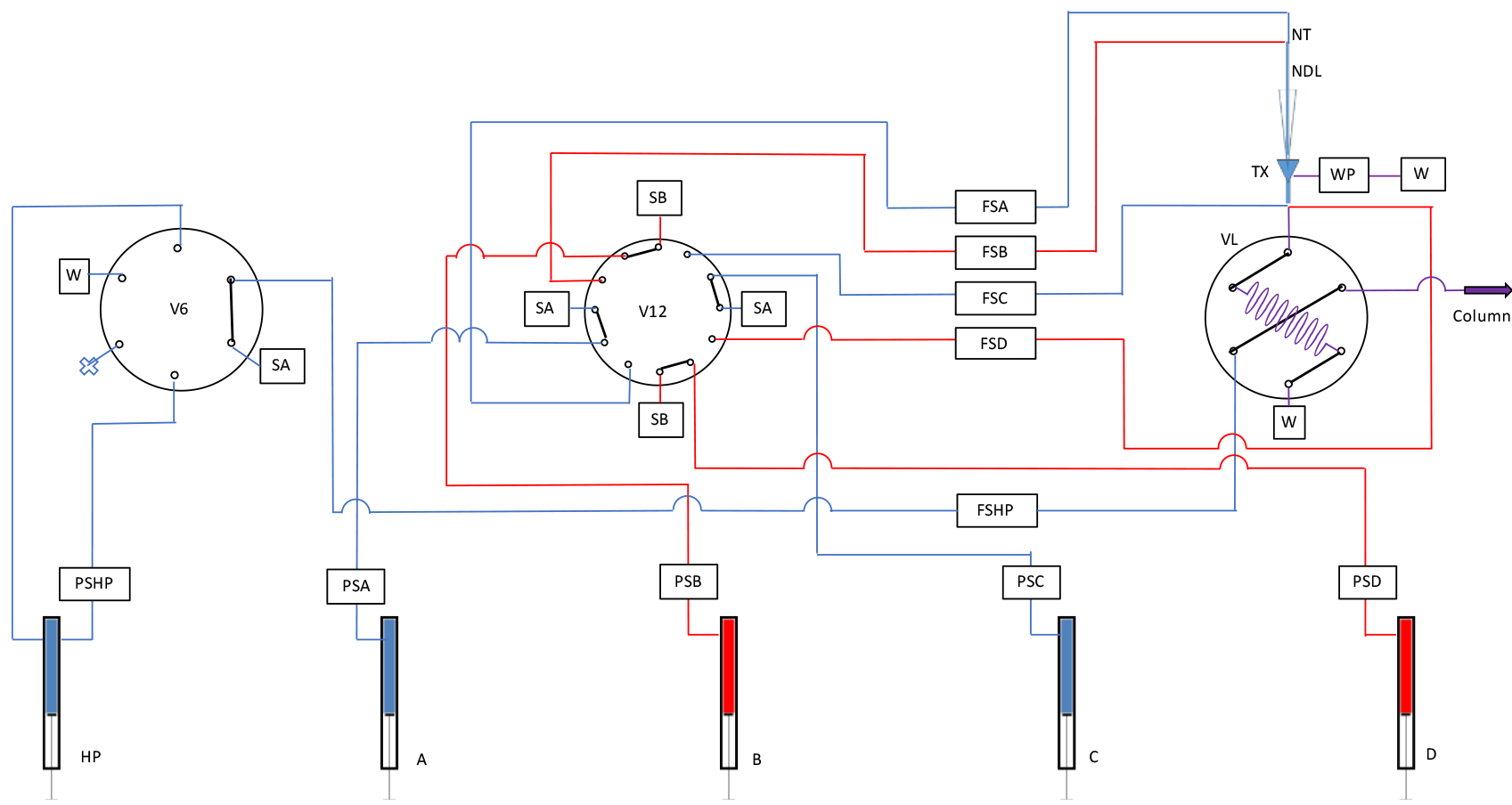
Evosep One flow diagram. Almost all of the system runs at low pressure (10-20 bar), increasing the lifetime and robustness of the LC. Only a single pump and flow path operates at high pressure and this does not involve any solvent mixing. HP: High-pressure pump, A/B/C/D: Low-pressure pumps, PSx: Pressure sensor x, FSx: Flow sensor x, WP: Waste pump, Sx: Solvent bottle x, W: Waste, V6: 6-port high-pressure solvent valve, V12: 12-port low-pressure solvent valve, VL: Loop valve. NT: Needle tee, NDL: Needle, TX: Tip cross.

The only common flow path is a storage loop, which is either connected to the low-or high-pressure sub-system and is controlled by a 6-port rotary valve (VL), see Figure 1. In this way, the high-pressure sub-system is always connected to the analytical or separation column but is either in-line or bypasses the storage loop. In contrast, the low-pressure sub-system is always connected to waste but either in-line or bypassing the storage loop. Thus, the storage loop becomes the bridge between the low-and high-pressure sub-systems.

The LC has an xyz manipulator, which picks up an individual disposable trap column (EvoTips) with its ceramic needle (NDL) and positions it in-line with the solvent flow path at the tip-cross (TX).

Pumps A and B then form a primary gradient that flows through the disposable trap column, eluting the analytes of interest. The organic content of this initial gradient is limited to less than 35% to ensure that only peptides of interest are eluted off the tips while unwanted compounds such as polymers, lipids, and other highly hydrophobic compounds remain bound to the single-use, disposable tips along with any particulate matter from the loaded samples. Furthermore, the final elution volume of this initial gradient is limited to few μl in order to ensure very precise elution and minimize bleeding of the more hydrophobic molecules. This “partial elution” concept will be further described in RESULTS AND DISCUSSION.

The two additional low-pressure pumps, C and D then modify the eluent to create an “offset” to the initial gradient. This has the purpose of lowering the organic contents, such that the analytes are initially retained on the analytical column. The offset gradient with the embedded analytes is moved into the storage loop before being switched in-line with the high-pressure pump, by means of which it is pushed towards the analytical column for high performance separation.

*Cell culture* - HeLa cells were cultured in high glucose DMEM with 10% fetal bovine serum and 1% penicillin–streptomycin (Life Technologies, Inc.). Cells were counted using an Invitrogen countess cell counter and stored after snap freezing at −80°C.

*Tryptophan fluorescence emission assay for protein quantification –* Protein concentrations were determined in 8 M urea by tryptophan fluorescence emission at 350 nm, using an excitation wavelength of 295 nm. Tryptophan at a concentration of 0.1 μg/μl in 8 M urea was used to establish a standard calibration curve (0−4 μl). We estimated that 0.1 μg/μl tryptophan are equivalent to the emission of 7 μg/μl of human protein extract, assuming that tryptophan on average accounts for 1.3% of human protein amino acid composition.

*Protein digestion* - For sample preparation we used the iST kit for proteomic samples (Kulak et al., 2014), starting with 106 HeLa cells according to the manufacturer’s instructions (P.O. 00001, PreOmics GmbH).

*Robustness optimization* - To test and optimize robustness, we injected and analyzed over 2000 times tryptic peptides of HeLa cells, initially in exploratory batches. For this experiment, we used the breadboard model of the Evosep One coupled to a LTQ Orbitrap instrument. All issues were protocolled and the system was optimized during the test and in the exploratory phase, the instrument was only stopped for the optimization of hardware and software components. The last 1500 HeLa samples were analyzed on a single column to analyze variation in the system and the wear of the column.

*Plasma proteomics* - Blood was taken by venipuncture using a commercially available winged infusion set and collection tubes containing EDTA and centrifuged for 15 min at 2000 g to harvest plasma. Blood was sampled from a healthy donor, who provided written informed consent, with prior approval of the ethics committee of the Max Planck Society. The plasma was distributed into a 96 well plate and subsequently processed with an automated sample preparation for Plasma Proteome Profiling as described previously (Geyer et al. 2016a).

*High throughput of low complexity samples* - the “UPS1 Proteomic Standard” (Sigma-Aldrich) was digested as indicated above using the PreOmics iST kit and the peptides were analyzed with the 200 samples/day method (5.6 min gradient) with a 2μl/min flow on a 5 cm C18 column (3μm particle size).

*Pre-fractionation* - Peptides for deep proteome analysis were fractionated using a reversed-phase Acquity CSH C18 1.7 μm 1 × 150 mm column (Waters, Milford, MA) on an Ultimate 3000 high-pressure liquid chromatography (HPLC) system (Dionex, Sunnyvale, CA) operating at 30 μl/min. Buffer A (5 mM ammonium bicarbonate) and buffer B (100% ACN) were used. Peptides were separated by a linear gradient from 5% B to 35% B in 55 min, followed by a linear increase to 70% B in 8 min. In total, 46 fractions were collected without concatenation. For nano-flow LC-MS/MS, the loading amount was kept constant at 500 ng per injection for the Easy-nLC 1200, while 500 ng from each fraction was loaded on an EvoTip.

*UV gradient storage experiment* - To assess the effect of diffusion as a function of storage time in a storage loop, we built a test rig to mimic Evosep One operation as illustrated in figure 2. A set of Zirconium nano pumps (Prolab Instruments, GmbH, Switzerland, pump A: 0.1% formic acid (FA) in H_2_O, Pump B: 0.1% FA, 1% acetone in acetonitrile) were programed to create the following composition profile: 0–5 min 5% B, 5–10 min 5–95% B, 10–13 min 95% B, 13–15 min 95–5%, 15–18 min 5%, 18–23 min 5–95% B, 23–25 min 95% B, 25–27 min 95–5% B, 27–30 min 5% B. This was delivered into a coiled (diameter 10 cm) fused silica storage loop (length 7 m, i.d. 100 μm, OD375, Polymicro Technologies). After a specified storage time had passed, a third Zirconium pump pushed the content out of the loop at a flow rate of 2 μl/min towards a UV detector (SpectraFlow 501, SunChrom) equipped with a nano-flow cell (5 nl) set to record the absorption at 265 nm. The storage loop and the three pumps were all connected to a standard 6-port Vici valve (Valco Instruments Co. Inc.) to control the flow path using a script.

**Fig. 2:**
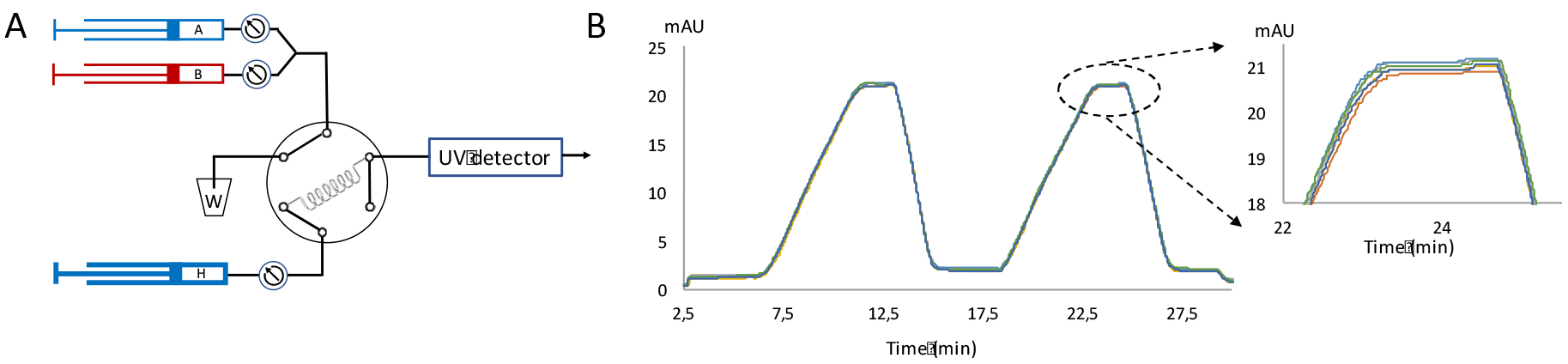
UV set up to test gradient storage. *A*, Flow diagram for testing potential gradient mixing during storage in the capillary loop. *B*, Profiles of the acetonitrile and water plugs that were recorded by the UV detector for different storage times. Profiles were almost completely superimposed, consistent with minimal mixing of the two phases during storage.

*Loading of EvoTips –* Tips were activated with consecutive 100 μl wash steps of 100% ACN, 50% ACN in 0.5% formic acid in H_2_O followed by two times 0.5% formic acid in H_2_O. BSA or HeLa peptides were loaded in 0.5% formic acid in H_2_O. The tip activation protocol was later optimized to use 2-propanol for wetting the C18 material prior to equilibration.

*High-pressure liquid chromatography and mass spectrometry* - LC-MS instrumentation consisted of a breadboard Evosep One coupled to an LTQ Orbitrap for the more than 2000 HeLa injection experiment, and the Evosep One production version coupled to an Q Exactive HF-X Orbitrap (Thermo Fisher Scientific) for all other experiments. Purified peptides were separated on the HPLC columns with 3 μm Reprosil-Pur C18 beads (Dr. Maisch, Ammerbuch, Germany) and dimensions indicated below in Table 1. On the LTQ Orbitrap MS, data were acquired with a Top6 data dependent shotgun method and with a Top12 method for the Q Exactive HF-X instrument. On the Q Exactive HF-X Orbitrap, the target value for the full scan MS spectra was 3 × 106 charges in the 300–1650 m/z range with a maximum injection time of 50 ms and a resolution of 60,000 at m/z 200. Fragmentation of precursor ions was performed by higher-energy C-trap dissociation (HCD) with a normalized collision energy of 27 eV (Olsen et al., 2007). MS/MS scans were performed at a resolution of 15,000 at m/z 200 with an ion target value of 5 × 104 and a maximum injection time of 25 ms. Dynamic exclusion was set to 15 s to avoid repeated sequencing of identical peptides.

*Deep proteome and DIA experiments* - HeLa cells were harvested at ~80% confluence by washing twice with PBS and subsequently adding boiling lysis buffer (6 M guanidinium hydrochloride (GndCl), 5 mM tris(2-carboxyethyl)phosphine, 10 mM chloroacetamide, 100 mM Tris pH 8.5) directly to the plate. The cell lysate was collected by scraping the plate and boiled for an additional 10 min, followed by micro tip probe sonication (Vibra-Cell VCX130, Sonics, Newton, CT) for 2 min with pulses of 1 s on and 1 s off at 80% amplitude. Protein concentration was estimated by Bradford assay, and the lysate was digested with LysC (Wako) in an enzyme/protein ratio of 1:100 (w/w) for 1 h, followed by dilution with 25 mM Tris, pH 8.5, to 2 M GndCl and further digested overnight with trypsin (1:100 w/w). Protease activity was quenched by acidification with trifluoroacetic acid (TFA) to a final concentration of ~1%, and the resulting peptide mixture was concentrated on Sep-Pak (C18 Classic Cartridge, Waters, Milford, MA). Elution was done with 2 ml of 40% acetonitrile (ACN), followed by 2 ml of 60% ACN. The eluates were combined and volume reduced by SpeedVac (Eppendorf, Germany), and the final peptide concentration was estimated by measuring absorbance at 280 nm on a NanoDrop spectrophotometer (NanoDrop 2000C, Thermo Fisher Scientific, Germany). For DIA samples, iRT peptides (Biognosys AB, Schlieren, Switzerland) were added prior to MS analysis according to the manufacturer’s protocol. For samples analyzed on the Evosep One, an in-house packed 12 cm, 150 μm i.d. capillary column with 1.9 μm Reprosil-Pur C18 beads (Dr. Maisch, Ammerbuch, Germany) was used, while samples analyzed on the Easy-nLC 1200 were separated in an in-house packed 15 cm, 75 μm i.d. capillary column with the specifications as described above. The column temperature was maintained at 40 °C using an integrated column oven (PRSO-V1, Sonation, Biberach, Germany) and interfaced online with the mass spectrometer.

*Data analysis* - MS raw files were analyzed by the MaxQuant software (Cox and Mann, 2008) and fragments lists were searched against the human Uniprot Reference Proteome without isoforms (April 2017 release with 21,042 protein sequences) by the Andromeda search engine (Cox et al., 2011) with cysteine carbamidomethylation as a fixed modification and N-terminal acetylation and methionine oxidations as variable modifications. The experiment for the 200 samples/day method was analyzed with the UPS1 FASTA file, downloaded from the homepage of Sigma-Aldrich (April 2018). We set the false discovery rate (FDR) to 0.01 at the peptide and protein levels and specified a minimum length of 7 amino acids for peptides. Enzyme specificity was set as C-terminal to arginine and lysine as expected using trypsin and LysC as proteases, and a maximum of two missed cleavages. An initial precursor mass deviation up to 7 ppm and a fragment mass deviation of 20 ppm were specified.

Data independent analysis (DIA) results were processed with Spectronaut version (11.0.15038.19.19667) (Biognosys, Zurich, Switzerland). A project specific spectral library was imported from the separate MaxQuant analysis of the combined analysis of the 46 pre-fractionated HeLa fractions, and DIA files were analyzed using default settings.

All bioinformatics analyses were done with the Perseus software (Tyanova, 2016) of the MaxQuant computational platform.

## RESULTS AND DISCUSSION

***Principle of analyte embedding in pre-formed gradients*** - Our key idea in making nano-LC as robust as high-flow LC was to decouple gradient formation from the high resolution, high-pressure separation on an analytical column. As in established peptide purification strategies, the peptides are first loaded on EvoTips (a form of solid phase extraction tips similar to StageTips (Rappsilber et al., 2003)). However, instead of eluting the peptides from the tips, drying them to remove the organic content and re-suspending them in injection buffer, we directly elute from the EvoTip into the capillary loop. This is accomplished at pressures of only a few bar by two syringe pumps A and B at flow rates of 10 to 20 μl/min (Fig 1). Note that an entire gradient can be stored in a several meters long fused silica capillary - already containing the individual peptides at the organic content where they elute from the C18 material. For instance, a 4 m long capillary of 100 μm inner diameter (i.d.) has a volume of 31.5 μl, sufficient for a subsequent analytical column separation of 31.5 min at 1 μl/min or 90 min at 350 nl/min.

We first asked if the gradient would be affected over time in the storage loop due to diffusion (Davis et al., 1995). Considering the very high aspect ratio of column length compared to i.d. (40,000 in the example above), this appears to be unlikely. Furthermore, in a similar capillary storage scheme in the RePlay system we did not observe such mixing (Waanders et al., 2008). To experimentally investigate this question, we placed defined plugs of ACN/1% acetone and water in the capillary loop, stored them for 0 or 60 min and monitored them with a UV detector (Fig. 2A, EXPERIMENTAL PROCEDURES). This did not lead to detectable mixing (Fig. 2B), confirming that storage of pre-formed gradients in a capillary loop is suitable for our purposes.

Having established that an analyte-containing gradient can be formed easily and be stored in a loop, the next challenge was to obtain high chromatographic resolution with the help of an analytical column. A common issue in pre-column setups is peak broadening because peptides eluting from the pre-column are not sufficiently retained on the analytical column. To solve this issue, and to take account of the relatively large elution volume from the EvoTip, we designed a gradient offset strategy. Once the EvoTip is sealed in-line with the solvent system, a gradient from pumps A and B subsequently elutes the peptides from the tip. Directly after the EvoTip, a secondary gradient from pumps C and D modifies the composition of the initial gradient and thus, reduce the effective organic content (Fig 3A,B). With the offset gradient, peptides eluting from the loop are shortly retained at the head of the column and thereby focused (Fig 3C). After separating on the analytical column, this results in the highest possible peak capacity. Note that due to the pre-formed and offset gradient the analytes are effectively loaded on the column in a sequential manner. Consequently, only a few percent of total peptide load is on the column at any given time (for instance, with a loop of 30 μl, a maximum of 3% for a 12 cm, 75 μm i.d. column which has a bed volume of less than one 1 μl).

**Fig. 3:**
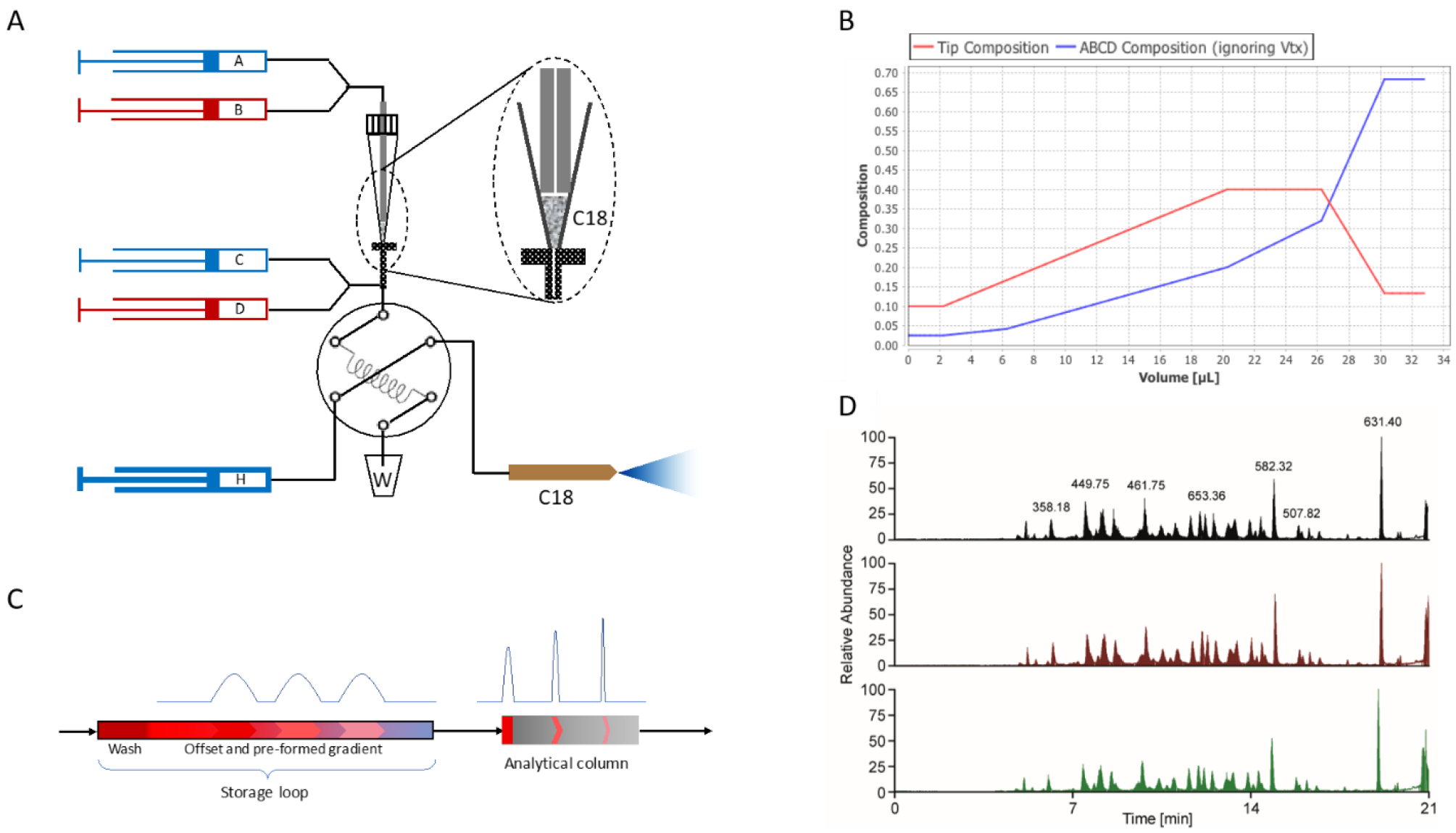
Pre-formed gradient. *A*, Peptides are eluted from the C18 containing EvoTip by pumps A and B. Pumps C and D, which are also low-pressure, form the final gradient, which is stored in the capillary loop together with the analytes. Subsequently, the valve switches and the high-pressure pump (H) simply pushes the gradient with its peptides over the analytical column. *B*, Composition of the gradient resulting from the confluence of the flows from pumps A, B and pumps C, D (x-axis designates the volume entering the storage loop). The proportion of acetonitrile is indicated on the y-axis. *C*, Analytes embedded in the storage loop are represented in red and as peak intensities. Due to the offset provided by pumps C and D, peptides are shortly retained at the head of the analytical column and elute with sharp peak widths. *D*, Comparison of three BSA chromatograms, demonstrating excellent reproducibility.

After generation of the gradient, the loop-valve switches the storage loop in-line with the high-pressure pump and the analytical column (Fig. 3A). The high-pressure pump then pushes the pre-formed and offset gradient with embedded, pre-separated peptides over the analytical column. The fact that almost all of the system’s functionality is contained in the low-pressure sub-system (Fig 1), should ensure long lifetime of the mechanical components, and opens up for ultra-precise flow manipulation, at a low risk of critical leaks and malfunction.

To test the Evosep One separation scheme, we loaded a BSA digest on an EvoTip and eluted it in a 21 min gradient from an 8 cm analytical column (100 μm i.d., 3 μm C18 beads). This resulted in low peak widths (4.8 s median FWHM) and corresponding column capacities. Multiple injections illustrate that the chromatograms are virtually superimposable (Fig 3D). An interesting consequence of our design is that it almost eliminates the loading and washing steps that are otherwise necessary between injections. Instead, the washing step is also encoded in the loop composition, and all remaining procedures take less than three minutes. This brings the total analysis time (injection to injection) very close to actual gradient time (21 min+3 min). Compared to conventional designs, this dramatically increases throughput, especially for short gradients, while avoiding the complexity and reproducibility issues of double column designs (Hosp et al., 2015).

***Robustness development and stress test*** - Having established the basic principles of operation, we constructed a breadboard model that incorporates all functional components. As far as possible, we chose industry leading standard components, such as the CTC Analytics auto sampler and Vici rotary valves, whereas other components were custom designed for our throughput and robustness requirements (EXPERIMENTAL PROCEDURES). Pump firmware development was done in house but for other software development, we used the Chronos environment, an industry standard and widely used platform, with a view to integrate our instrument with the different MS manufacturers.

To fine-tune operation and optimize robustness, we injected 1 μg of a tryptic HeLa cell digest over 2000 times in a consecutive manner. We logged all issues over time and stopped the test only to optimize hardware or software components. A substantial percentage of the runs within the first 250 samples suffered from sample loss. We were able to trace this to an imperfect seal of the autosampler needle and the tip. In a first step, we optimized the needle, which resulted in an immediate reduction of errors. After changing the seal between the tip and the entrance of the flow path as well, these issues were completely eliminated (Fig 3A,4A). From injection 513 on, all instrument related issues appeared to be resolved. We then mounted a new column to test the new “partial elution” concept (as described in EXPERIMENTAL PROCEDURES) in subsequent injections. Over these 1500 samples and the total ion current remained unchanged until the end of the experiment (Fig 4B). A few LC-MS runs were blank, but this turned out to be due to incorrect manual loading of the corresponding EvoTips.

**Fig. 4:**
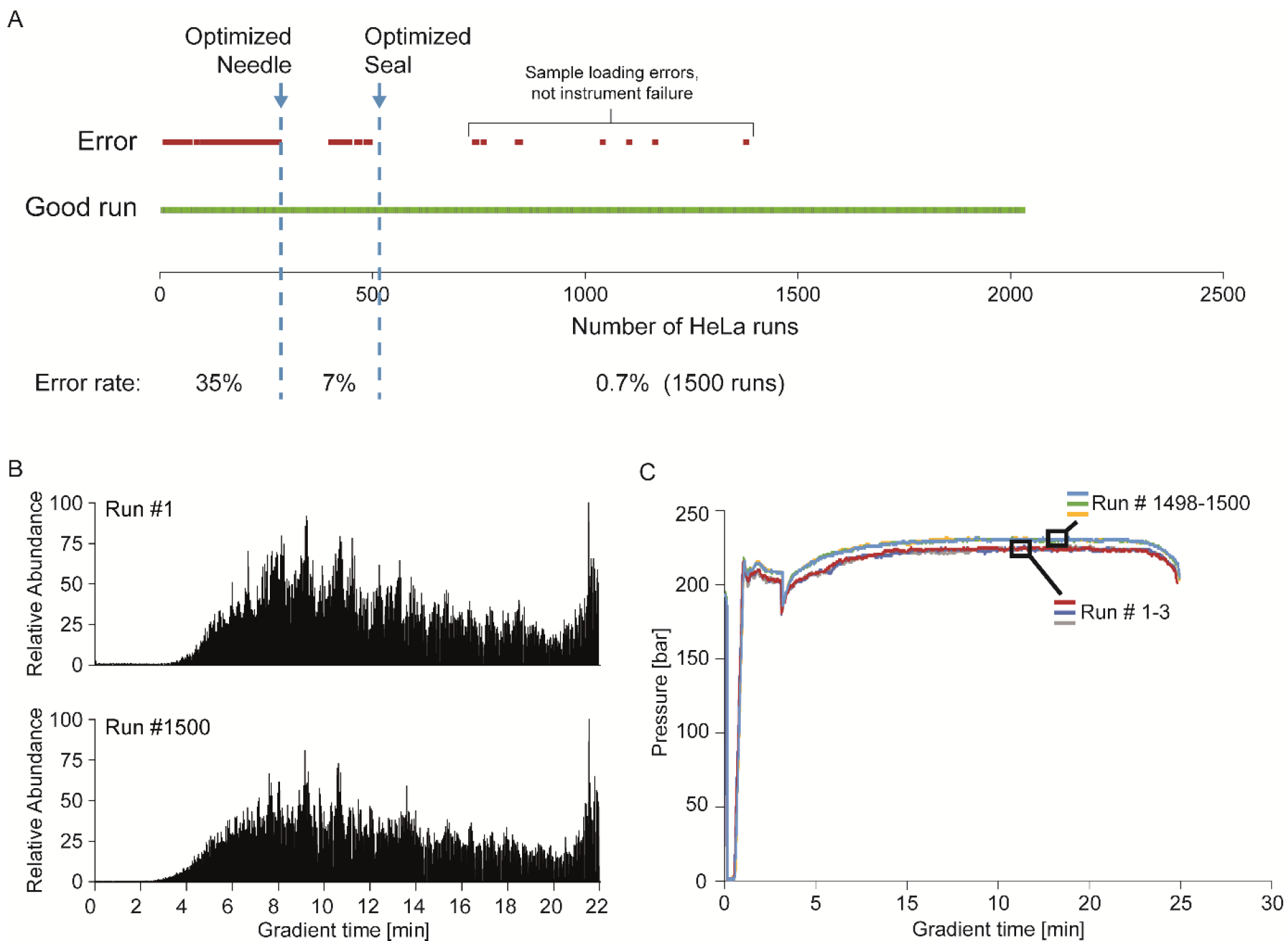
Robustness evaluation. *A*, Error frequency during the development phase of the system assessed by consecutive measurements of HeLa digests. *B*, The first and last chromatogram of a HeLa digest in a series of 1500 measurements. *C*, Pressure profiles over the gradient for the first and last three HeLa digest of the same experiment.

We also recorded the pressure profiles for all the runs. Validating the partial elution concept, there was only a very slight increase in backpressure, indicating that the column had remained free of deposits and as further evidence of the effect, the TICs of runs 1 and 1500 were indeed highly similar and showing no decay in separation performance of the column. Pressure profiles of adjacent runs were virtually indistinguishable (Fig 4C) The Evosep One was intended and constructed for high throughput applications, with a particular focus clinical analysis. Blood plasma is the most widely analyzed clinical matrix, with millions of samples drawn daily. Yet it is difficult to analyze plasma robustly by nano-LC MS, mainly because of the large number of non-protein blood components. To demonstrate clinical applicability of the system, we employed our automated sample preparation pipeline - termed Plasma Proteome Profiling (Geyer et al. 2016a). Plasma samples were prepared and loaded on the EvoTips in a 96 well format, using a robotic platform. The total measurement time for the 96 samples on the Evosep One was less than two days, corresponding to a throughput of 60 samples per day. Reproducibility over all 96 independent, parallel sample preparations and injections of the same original plasma was excellent (median Pearson correlation coefficient of 0.98) over all runs (Fig. 5B). For clinical decision making based on the concentration of biomarkers, it is crucial to ensure low carry-over from one analysis to the next. Therefore, we performed a cross contamination experiment with six alternating injections of plasma and blanks (Fig. 5B). The average carry-over was as low as 0.07% and 80% of this can be traced back to just 20 peptides (Fig. 5C).

**Fig. 5:**
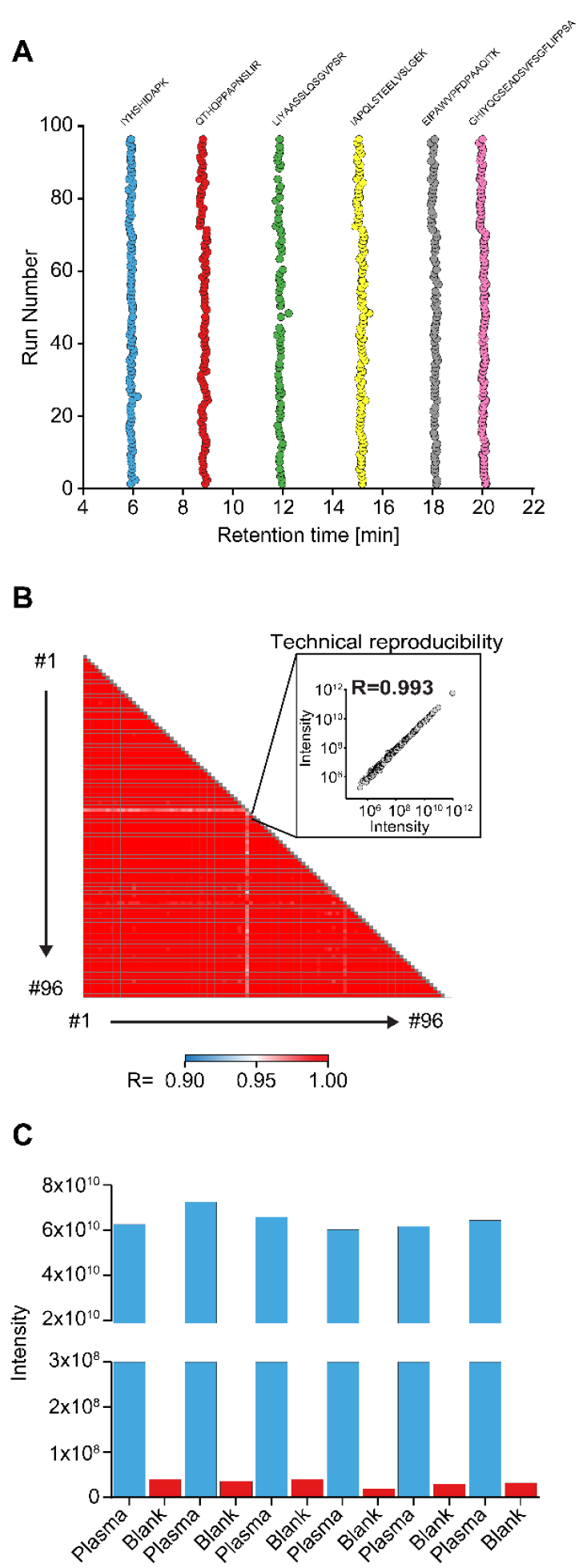
Clinical applicability to the plasma proteome. *A*, Retention time stability of selected peptides spanning a range of elution times over 96 plasma proteome runs. *B*, Pearson correlation matrix comparing all 96 plasma runs to each other. A single correlation graph with the median Person value is shown in the inset. *C*, Summed total peptide intensities in alternating plasma and blank runs.

***Design of methods for desired throughput and depth*** - Based on the principles explained above and the experiences from the robustness testing on the breadboard model, we then constructed the production unit. We devised a number of standard gradients and column combinations tailored to different applications, ranging from high throughput quality control of low complexity samples, through comprehensive proteomics using fractionation, to the in depth single run characterization of complex proteomes. In particular, the short gradients made possible by the Evosep system can be used for low complex samples and the longer gradients for more complex samples. Table 1 contains the optimized gradients and column dimensions for the diverse use cases and sample types.

**Table 1:** Evosep One methods. For ease of use, five pre-set methods have been optimized to provide the best performance to time compromise. They are defined by the total number of samples that can be run per day rather than referring to the length of the gradient.

| Throughput | Cycle time | Gradient length | Overhead | Flow rate | Column length / ID |
| --- | --- | --- | --- | --- | --- |
| Samples / day | [min] | [min] | [min] | [μl / min] | [cm] / [μm] |
| 300 | 4.8 | 3.2 | 1.6 | 3.0 | 4 / 150 |
| 200 | 7.2 | 5.6 | 1.6 | 2.0 | 4 / 150 |
| 100 | 14.4 | 11.5 | 2.9 | 1.5 | 8 / 100 |
| 60 | 24.0 | 21.0 | 3.0 | 1.0 | 8 / 100 |
| 30 | 48.0 | 44.0 | 4.0 | 0.5 | 15 / 100 |

We first wished to demonstrate the possible throughput on low complexity sample. We digested the “UPS1 Proteomic Standard” (EXPERIMENTAL PROCEDURES) and used the 5.6 min gradient with the 2 μl/min flow on the 5 cm column (200 samples/day method). In one day, this indeed resulted in 200 data sets with very consistent protein coverage (Fig. 6). The UPS1 should contain 48 different proteins but curiously four of them were never identified. As this standard is equimolar this is not an issue of dynamic range. Furthermore, the remaining 44 proteins were quantified essentially completely in all runs (average of 43.5±1) (Fig. 6). We conclude that the remaining proteins were likely missing from the kit. The high throughput for low complexity samples would be very interesting for single protein identification experiments in gel bands, for instance, or for contaminant analysis in recombinant protein expression in biotechnology. In many cases, it could also be sufficient for somewhat more complex mixtures such as those resulting from pull-down experiments.

**Fig. 6:**
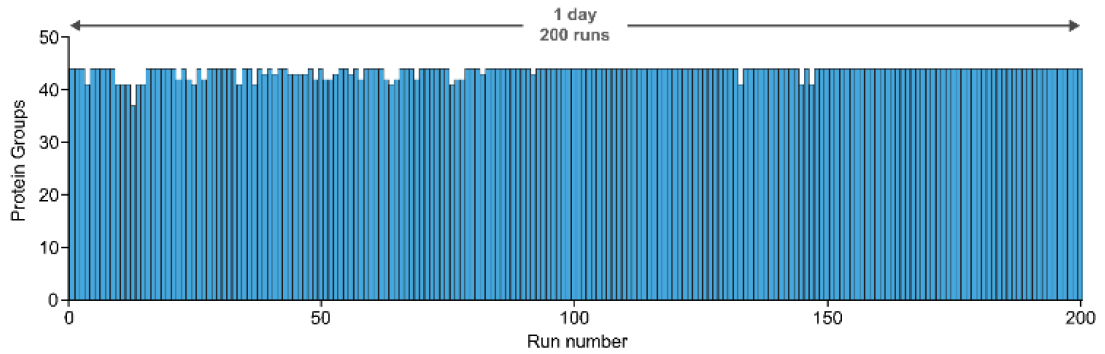
High throughput for low complexity samples. Technical replicates of a digest of the UPS1 Proteomic Standard were injected 200 times with the 200 samples/day method of Table 1. The number of identified proteins for each sample is shown as a bar graph in chronological order.

***Rapid generation of in-depth mammalian cell line proteomes*** - Having shown the applicability of the system for low complexity sample in high throughput, we next investigated the rapid characterization of fractionated, high complexity proteomics samples. A fractionation step is very common in the analysis of cell line or tissue proteomes, but usually comes with the caveat of a drastic increase in measuring time as the number of factions increases.

We built on a recently described strategy that combined extensive high pH reversed-phase peptide pre-separation in a first HPLC dimension without ‘concatenation’ of the resulting fractions (Bekker-Jensen et al., 2017). Up to 70 such fractions were analyzed in relatively short gradients of 30 min, allowing for overall high peptide loading and high combined peak capacity and making optimal use of the high acquisition speed of state of the art mass spectrometers (Kelstrup et al., 2014). This resulted in a very deep coverage of cell line and tissue proteomes, on par with RNA-seq results (Bekker-Jensen et al., 2017). A bottleneck of the workflow was the relatively low utilization of the mass spectrometer, due to the washing, equilibration and loading times of the HPLC, which are minimized with the Evosep system.

To characterize the efficiency for fractionated proteomes and to compare this to the Easy-nLC 1200 used as a standard in our laboratories as well as in the study described above, we performed an analysis of 46 HeLa fractions on both systems. Each of the fractions was divided and separately measured on the Easy-nLC and the Evosep One on the same MS instrument, recording, total instrument time, the time utilized for gradients and the numbers of peptides and proteins identified. The Easy-nLC 1200 was run with our previously optimized 15 min gradients, whereas we used the 21 min gradient of the 60 proteome/day method for the Evosep One.

As expected due to the short overhead time between runs, the Evosep One was significantly more efficient in terms of utilization of the mass spectrometer. A full 88% of the total analysis time of 18.4h was spent on data acquisition (Fig. 7A). In contrast, the Easy-nLC 1200 occupied the mass spectrometer for 28.3 h, but only 14.6 h (52%) were productively used. This difference did not come at the expense of the numbers of identified peptides and proteins, which was very comparable with 132,850 peptides (9918 proteins) and 130,450 peptides (9603 proteins) for the Evosep One and the Easy-nLC 1200, respectively. A detailed view of peptides identified in each fraction separately or cumulatively, showed that they are very similar (Fig. 7B,C). This confirms our conclusion that the design principle of the Evosep One resulted in saving substantial measurement time (35% in this case), at undiminished performance. For longer gradients, the proportional time savings would be lower, however, given the high price of modern mass spectrometers, they would still be economically attractive. The above experiments show that the Evosep is well suited for the in depth characterization of proteomes via the rapid analysis of the high pH or other fractions that are commonly used in proteomes. While we employed label-free quantitation here, the results should equally apply to isobaric labeling strategies. We also note that an average of 2700 proteins were identified in these fractions. There are a number of proteomics strategies that produce many fractions, such as thermal shift assays (Savitski et al., 2014) or organellar proteomics (Itzhak et al., 2016), and our approach opens up for strategies to rapidly and robustly measure these.

**Fig. 7:**
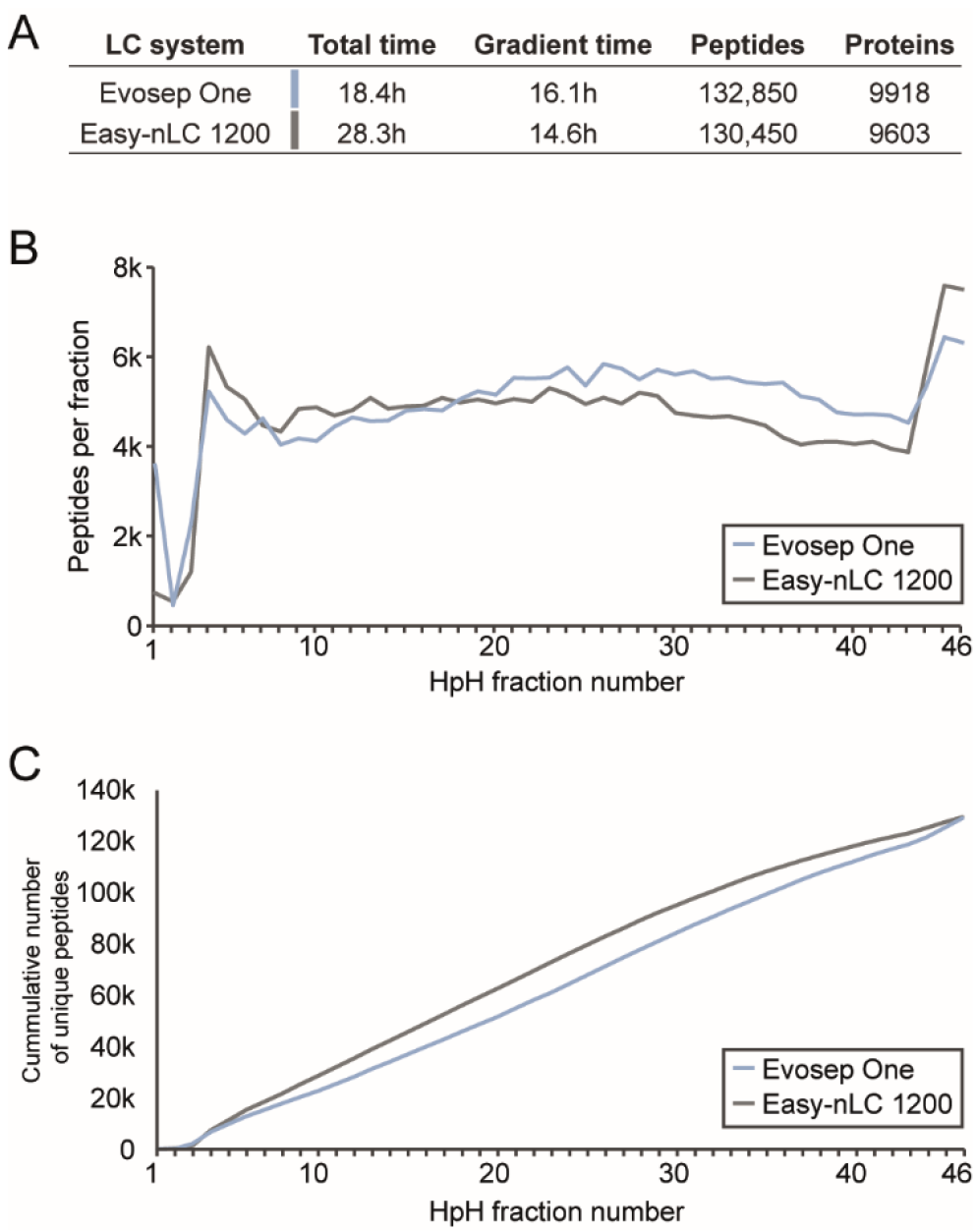
Rapid generation of mammalian cell line proteomes. *A*, Table for the comparison of the Evosep One with the Easy-nLC 1200, including total measurement time, gradient time and the numbers for identified proteins and peptides. *B*, Numbers of identified peptides per fraction over the 46 high pH reversed-phase fractions for both LC systems. *C*, Cumulative numbers of unique peptides across the fractions.

**Fig. 8:**
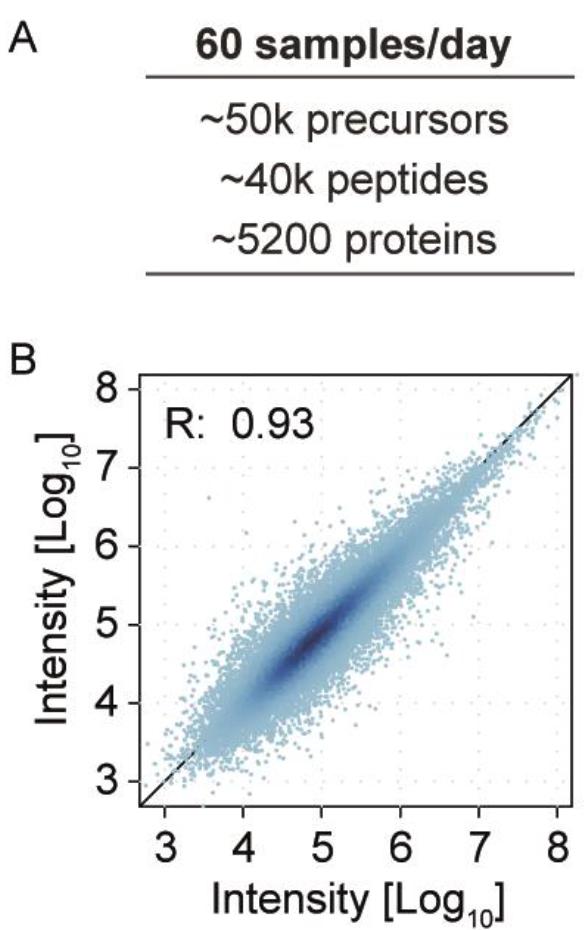
Rapid generation of mammalian cell line proteomes. *A*,Average number of precursors, identified peptides and protein groups for DIA measurements with 21 min gradients on the Evosep One. *B*, Average Pearson correlations coefficient for the comparison of the precursor intensities of two replicate HeLa analysis.

***Single shot, high throughput HeLa proteomes using DIA*** - The experiments described so far all used data dependent acquisition (DDA). However, data independent acquisition (DIA) is becoming increasingly popular and competitive (Bruderer et al., 2017). In our hands, we have found DIA to perform particularly well with relatively short gradients on fast and high resolution Orbitrap analyzers (Kelstrup et al., 2018). The Evosep One with its fast turn-around between runs appeared to be a good addition to this strategy and we were curious to see how deep the proteome could be covered with such a combination. For this purpose, we made use of the very extensive peptide library generated in our previous experiments of the 46 fractions of HeLa digests using the Spectronaut software with one percent FDR at both precursor and protein levels. We applied a DIA method with a precursor mass range (m/z 361–1033), which covers the majority of tryptic HeLa peptides within 4 s cycle time. This was achieved by one full MS (128 ms transient) followed by 56x HCD-MS/MS spectra (64 ms transient) with 13 m/z isolation widths allowing 1 Th overlap between windows. This setup allowed us to analyze 60 HeLa proteomes in a single day. Given the short gradients, the proteome coverage was very high with an average of 5200 proteins from 40,000 peptide identifications. Note that this implies a very high matching efficiency, because 80% of the 50,000 precursors were statistically significantly identified. Measured over time, this enabled identification of 250 unique proteins per gradient minute throughout the gradient. Moreover, there was a high consistency in the number of identified protein groups with a high overlap between replicates of 92%. The comparisons of the individual runs showed an average Pearson correlation coefficients of R=0.93, which indicates robust quantification. This suggests that the Evosep system is also a well suited instrument for high-throughput DIA analysis.

## CONCLUSION

Despite the great technological advances in high sensitivity nano-flow MS-based proteomics, the robustness and throughput have been weak links even in state of the art MS-based proteomic workflows. This has led to a move towards microflow systems - especially with a view towards clinical applications - however, at the cost of sensitivity (Fu et al., 2018). Here, we have introduced an entirely novel concept based on the pre-formation of gradients at relatively high-flow and low-pressure. This pre-stored gradient already has the analytes embedded and is moved across a high-resolution column by a single, high-pressure pump. Based on these principles, we first designed a breadboard system that was progressively developed into a commercial HPLC system - the Evosep One. We established that pre-storing of the gradient, followed by ’sharpening of the peaks at the head of the analytical column, assures full chromatographic peak capacity of the overall system. Together with the EvoTip as a disposable sample clean up cartridge, the system is designed for sensitivity, throughput, and robustness - tailor made for large clinical studies. To test this, we performed thousands of runs with cell lysates as well as complex clinical samples such as blood plasma. We found that the decoupling of gradient formation with a low-pressure system and the high-pressure peptide separation ensured stable and uninterrupted operation without instrument related issues or deterioration in chromatographic performance. As expected from its design, the Evosep One proved to have minimal or absent cross contamination and very high consistency of label-free quantitation results across numerous injections.

The time required for the formation of the pre-stored gradient, including the washing step, happens very quickly, effectively eliminating idle time of the mass spectrometer between injections. This opens up for the rapid analysis of samples of medium complexity, as we demonstrated with the measurement of 200 standard mixtures (UPS1) in a single day. Deep proteomes are typically achieved after extensive fractionation. In this context, the fast turn-around of the Evosep One ensures very high utilization of the MS instrumentation as we show by the analysis of 46 HeLa fractions in 18 h. Finally, we used a state of the art data independent workflow that enabled a remarkable depth of proteome depth of 5,200 proteins in only 21 min (60 samples/day method). With ongoing developments on the mass spectrometric side, the proteome depth is likely to improve further. The ability to analyse 60 proteomes in a single day opens up entirely new possibilities in proteomics in general and in clinical proteomics in particular.

## ACKNOWLEDGEMENT

We thank all members of the department of Proteomics and Signal Transduction at the Max Planck Institute of Biochemistry in Martinsried for help and discussions, and Gaby Sowa for technical assistance.

